# Lipoxin B_4_ Mitigates TRPV4-Activated Müller Cell Gliosis During Ocular Hypertension

**DOI:** 10.1101/2025.09.17.673801

**Authors:** Matangi Kumar, Shruthi Karnam, Shubham Maurya, Rama Nagireddy, John G. Flanagan, Karsten Gronert

## Abstract

**Purpose:** Müller glia play dual roles in glaucoma, contributing to both retinal homeostasis and neuroinflammation; their activation by elevated intraocular pressure through the mechanosensitive channel TRPV4 promotes a reactive state that drives retinal ganglion cell (RGC) loss. Lipoxin B_4_ (LXB_4_), an endogenous lipid mediator produced by retinal astrocytes, has been shown to suppress glial reactivity and directly protect RGCs. This study investigated whether LXB_4_ modulates TRPV4-driven Müller glial activation and inflammation and whether Müller glia themselves contribute to this retinal lipoxin pathway.

**Methods:** Ocular hypertension (OHT) was induced in mice via a silicone oil model, and reactive Müller glia were isolated via magnetic sorting for transcriptomic analysis. *In vitro*, primary and immortalized Müller glia were treated with a TRPV4 agonist with or without LXB_4_. Glial reactivity was assessed by flow cytometry, immunostaining, qPCR, and western blotting. LC–MS/MS-based lipidomics was used to quantify lipoxin pathway metabolites, and single-cell RNA-seq was used to examine transcriptional responses to LXB_4_ treatment. GFAP and TRPV4 expression was evaluated via immunohistochemistry in retinal sections.

**Results:** RNA bulk-sequencing analysis and qPCR revealed that Müller glia express both 5- and 15-lipoxygenase. Lipidomic analysis confirmed that the lipoxin pathway is functional and that Müller glia endogenously produce LXB_4_, establishing this essential cell type as a source of anti-inflammatory and neuroprotective LXB_4_ in the retina. TRPV4 activation induced a reactive glial phenotype characterized by increased GFAP and IL6 expression, increased STAT3 phosphorylation, and increased production of lipoxins, suggesting that biomechanical stress simultaneously triggers both gliosis and protective lipid signaling. Treatment with LXB_4_ suppressed TRPV4-induced gliosis *in vitro* by downregulating IL6 and inhibiting STAT3 activation, and *in vivo* by reducing the expression of *Stat3, Il6*, and *TNF-*α during OHT while attenuating TRPV4 upregulation in Müller glia.

**Conclusion:** Müller glia are a significant source of LXB_4_ in the retina. This neuroprotective Müller glia pathway is amplified during chronic TRPV4 activation to counter-regulate gliosis. The findings support targeting of the TRPV4–lipoxin pathway as a potential approach to protect against OHT-induced neurodegeneration in glaucoma.

## Introduction

Glaucoma, a group of optic neuropathies, is characterized by the progressive degeneration of retinal ganglion cells (RGCs) and their axons. Elevated intraocular pressure (IOP), a major risk factor, disrupts retinal homeostasis and activates inflammatory pathways that accelerate RGC loss [1,2]. While IOP-lowering therapies slow disease progression, they do not fully prevent RGC degeneration, underscoring the need for neuroprotective strategies that target underlying mechanisms. Emerging evidence suggests that glia–neuron interactions, including those involving Müller glia, play a critical role in glaucomatous neurodegeneration [3]. However, the molecular mechanisms underlying the contributions of glia to RGC loss remain unclear. Elucidating these pathways is essential for developing therapies that preserve RGCs and slow disease progression.

Müller glia are the principal macroglial cells of the retina and extend from the inner limiting membrane to the outer limiting membrane [3,4]. This unique architecture enables them to detect and respond to mechanical, metabolic, and inflammatory stress across all retinal layers [5]. Under physiological conditions, Müller glia regulate osmotic balance, support neuronal metabolism, and maintain ion and water homeostasis [3,6]. Following retinal injury, these cells become reactive, characterized by glial fibrillary acidic protein (GFAP) upregulation [3]. Chronic gliosis can disrupt tissue repair, promote inflammation, and accelerate RGC degeneration [5,7].

A key pathway driving Müller glial reactivity involves the nonselective, mechanosensitive cation channel transient receptor potential vanilloid 4 (TRPV4) [8,9]. TRPV4 is expressed in both RGCs and Müller glia and responds to mechanical insults, such as those caused by elevated IOP, by facilitating calcium influx and triggering downstream inflammatory signaling [10,11]. Sustained TRPV4 activation in Müller glia induces a reactive phenotype characterized by proinflammatory cytokine release and upregulated GFAP expression [8,9,12]. This process involves Janus Kinases (JAK) and Signal Transducers and Activators of Transcription (STAT) signaling, including IL-6/gp130 activation, and direct STAT3 binding to the GFAP promoter, ultimately contributing to RGC apoptosis in glaucoma [13–15]. Pharmacological activation of TRPV4 in Müller glia recapitulates key features of IOP-induced insult, making it a robust *in vitro* model for glaucomatous neuroinflammation [8,9,12].

Lipoxins (LXA_4_ and LXB_4_) are endogenous eicosanoids whose canonical role is to counter-regulate chronic inflammation. They are synthesized through cellular interactions between selective cell types expressing 5-lipoxygenase (5-LOX) and 15-LOX [16–18]. The anti-inflammatory action are well established for LXA_4_ and LXA_4_ stable analogs and include inhibiting leukocyte recruitment, JAK/STAT signaling, and NF-κB–driven cytokine production [19–21]. LXB_4_’s mechanisms of action is distinct from LXA_4,_ not mediated by the LXA_4_ receptor and remains to be fully defined. We recently identified lipoxin B_4_ (LXB_4_) as an endogenous lipid mediator in the healthy retina and as a promising neuroprotective target to prevent RGC death in glaucoma models [22,23]. Its novel neuroprotective actions in the retina and optic nerve are more potent than those of LXA_4_ and restore the homeostatic function of RGC, astrocytes and microglia [23–25]. In the retina, astrocytes generate LXB_4_ as part of their homeostatic function [23]. Our recent findings also revealed that treatment with LXB_4_ reduces glial reactivity in both astrocytes and Müller glia during ocular OHT [24]. However, whether Müller glia can endogenously produce LXB_4_ or if LXB_4_ modulates TRPV4-induced reactivity remains unclear. Given Müller glia’s shared homeostatic roles with astrocytes and their abundance (accounting for ∼90% of all retinal macroglia), it is plausible that Müller glia serve as a major source of retinal LXB_4_ and/or their metabolic precursors [26].

In this study, we investigated whether Müller glia express lipoxygenases required for LXB_4_ biosynthesis (5-LOX and 15-LOX) and whether they generate LXB_4_. We also examined whether LXB_4_ treatment counterregulates TRPV4-induced Müller glial reactivity in both *in vitro* (TRPV4 agonist–treated Müller glia) and *in vivo* (mouse OHT) models. This study defines a novel glia–lipoxin regulatory mechanism and provides new insight into potential neuroprotective strategies for glaucoma.

## Materials and methods

### Mice

C57BL/6J mice were purchased from The Jackson Laboratory (Bar Harbor, ME). All animal procedures were approved by the Institutional Animal Care and Use Committee (IACUC) at the University of California, Berkeley. The mice were housed under a 12-h light/12-h dark cycle with *ad libitum* access to food and water.

### Mouse model of ocular hypertension

Silicone oil–induced OHT was induced in 8-wk-old mice via an established protocol [27–29]. The mice were anesthetized by intraperitoneal (*i.p.*) ketamine (100 mg/kg) and xylazine (10 mg/kg), and topical ocular anesthesia was achieved with 0.5% proparacaine hydrochloride (Sandoz, Princeton, NJ). Under a surgical microscope, pharmaceutical-grade silicone oil (Alcon, Fort Worth, TX) was injected into the anterior chambers of both eyes using a 33 G Hamilton glass syringe (Reno, NV), avoiding contact with the lens and corneal endothelium. Chronic mild OHT was induced by injecting 1.2 µL of silicone oil. Acute severe OHT was induced by injecting 1.8 µL of silicone oil. Sham-injected control mice received a corneal incision created with a sterile 31 G paracentesis needle without silicone oil. After each procedure, 0.3% tobramycin ophthalmic solution (Tobrex; Alcon, Fort Worth, TX) was applied topically. The IOP was measured via a rebound tonometer, as described previously [23,24]. Tissues from the chronic mild OHT group were collected by euthanizing mice at 2, 3, 6, and 8 wk post-injection. Eyes were enucleated and fixed in 4% paraformaldehyde (PFA) for immunohistochemistry (IHC). For the acute severe OHT group, mice were euthanized at 1 wk postinjection, and eyes were enucleated and fixed in 4% PFA for IHC.

### Quantitative PCR

Total RNA was isolated from mouse retinas and Müller glia cultures via TRIzol reagent (Invitrogen, Waltham, MA) according to the manufacturer’s instructions and quantified using our previously published protocol [24]. First-strand cDNA was synthesized from 1 µg of RNA via iScript Reverse Transcription Supermix (Bio-Rad, Hercules, CA). The transcripts for *Il6, Stat3, TNF*-α*, Alox5, and Alox15* were quantified via GoTaq PCR master mix (Promega, Madison, WI) in a OneStep Plus qPCR system (Applied Biosystems, Waltham, MA). *Gapdh* served as the control for normalization. The mouse primers used in this study are listed in Table 1. Primers for rat genes used in this study are listed in Table 2.

### Bulk RNA sequencing

#### Retinal Dissociation and Müller Glia Enrichment

Three weeks after silicone oil–induced OHT, the mice were euthanized, and neural retinas (n = 4 per pooled sample) were dissected free of retinal pigment epithelium. Retinal tissue was dissociated into single cells via a papain dissociation system (Worthington, Columbus, OH), passed through a 40 µm cell strainer, and washed with PBS containing 0.5% bovine serum albumin (BSA). Dead cells and debris were depleted with the Dead Cell Removal Kit (Miltenyi Biotech, Bergisch Gladbach, Germany). The remaining cell suspension was incubated on ice for 10 min with 15 µL of anti-GLAST (ACSA-1)-APC antibody at a 1:50 dilution (Miltenyi Biotech) protected from light. After washing, the cells were centrifuged (300 g, 5 min), resuspended in 90 µL of MACS buffer and 10 µL of anti-APC microbeads (Miltenyi Biotech), and then incubated at 4°C for 15 min. Following a wash, the cells were passed through an LS column (Miltenyi Biotech) in a QuadroMACS separator. Unlabeled (GLAST⁻) cells were collected as flow-through; the column was washed 3x with 3 mL of MACS buffer. GLAST⁺ Müller glia were eluted with 5 mL of MACS buffer via a plunger.

#### RNA isolation and library preparation

Total RNA from enriched Müller glia was extracted via the Arcturus™ PicoPure RNA Isolation Kit (Thermo Fisher, Waltham, MA). RNA quality was assessed via an Agilent 2100 Bioanalyzer (Agilent Technologies, Santa Clara, CA), and samples with an RNA integrity number (RIN) of 7.0 were used for library construction. cDNA was synthesized and amplified via the SMARTer v4 Ultra-Low Input RNA Kit (Clontech, Mountain View, CA). cDNA was fragmented on a Diagenode Bioruptor Pico (Diagenode, Denville, NJ), and sequencing libraries were prepared with the KAPA HyperPrep DNA Kit (Roche, Basel, Switzerland). Libraries were pooled and sequenced on an Illumina NovaSeq 6000 S4 flow cell (QB3 Genomics Core, UC Berkeley; RRID: SCR_022170) to generate 150 bp paired end reads.

#### Bioinformatic analysis

The raw base call files were demultiplexed via Illumina bcl2fastq2 (v2.20). Adapter trimming and quality filtering were performed with Trim Galore (v0.6.6). Quality metrics were assessed with FastQC (v0.11.9). Cleaned reads were aligned to the mouse genome (mm39, UCSC) via STAR (v2.7.1a). Gene counts were obtained via featureCounts (version 1.5.3). Downstream analysis was conducted via RStudio (version 4.2.0). Transcript abundance was expressed as transcripts per million (TPM) via the normalizeTPM() function to normalize transcripts for gene length and sequencing depth. Differential gene expression was determined with DESeq2 (p.adj<0.05, |Log_2_FC| > 1). Gene set enrichment analysis (GSEA) was performed with fgsea (p.adj<0.05, |Log_2_FC| > 1), and the results were visualized in ggplot2 [24].

### Cell culture

#### Primary Müller glia

Primary mouse Müller glia were isolated from postnatal day 8–10 C57BL/6J pups [30–32]. Retinas were dissociated with papain (Worthington, Columbus, OH) into a single-cell suspension. The cells were plated on 6-well plates coated with 0.1% gelatin in Dulbecco’s modified Eagle’s medium (DMEM) (Gibco, Waltham, MA) supplemented with 10% fetal bovine serum (FBS) (Avantor Seradigm; Radnor, PA) and 1% penicillin–streptomycin (Gibco, Waltham, MA). Cultures were maintained at 37°C in a humidified 5% CO₂ incubator. Once confluent, the cells were passaged onto fresh gelatin-coated plates. Müller glia identity was confirmed after 2 passages by immunofluorescence staining for Vimentin, Glutamine Synthetase, and Sox9. All the assays were conducted using cells after passage 3.

#### Immortalized rat Müller glia ( rMC-1) Müller glia

The immortalized rat Müller glia cell line (rMC-1; T0576) was obtained from the UC Berkeley Cell Culture Facility. The cells were cultured in DMEM supplemented with 10% FBS, 1% MEM NEAA (Gibco, Waltham, MA), 1% sodium pyruvate (Gibco, Waltham, MA), and 1% penicillin– streptomycin (Gibco, Waltham, MA). Immortalized cells were used in place of primary cells for experiments where abundant protein and RNA were needed. All experiments were performed using cells at passages 2-10. FBS ( Avantor Seradigm; Radnor, PA) was used for all *in vitro* work to control for potential biological variations between different batches.

#### In vitro treatments

Cells (both primary and immortalized) were treated with a TRPV4 agonist at a concentration of 10 µM GSK1016790A (Millipore Sigma, G0798) or with an equivalent volume of vehicle (DMSO, < 0.02%) in culture medium. For LXB_4_ treatment, cells were incubated in 1 µM LXB_4_ (Cayman Chemical, Ann Arbor, MI) or the corresponding vehicle (ethanol < 0.01%)). For cotreatment experiments, the cells were pretreated with LXB_4_ for 20 min before the addition of GSK1016790A.

### Immunofluorescence staining

Enucleated eyes were fixed overnight in 4% PFA at 4°C and then dehydrated in a sucrose gradient (10%, 20%, and 30% sucrose in PBS) until the tissues sank. Eyes were embedded in optimal cutting temperature (OCT) medium and frozen at −80°C for immunohistochemistry (IHC). Ten-micron sections were cut on a Leica CM1900 cryostat (Leica Microsystems, Wetzlar, Germany) and mounted on Superfrost Plus slides (Thermo Fisher, Waltham, MA). Immunocytochemistry (ICC) was performed on primary and immortalized Müller glia were grown on glass coverslips, washed with PBS, and fixed in 4% PFA for 15 min at room temperature (RT). The sections and coverslips were permeabilized and blocked for 1 h in PBS containing 10% donkey serum and 0.25% Triton X-100 at RT. Primary antibodies against GFAP (1:1000; Abcam, Cambridge, UK), vimentin (1:500; Abcam, Cambridge, UK), Sox9 (1:200; Abcam, Cambridge, UK), Iba-1 (1:200; Cell Signaling, Danvers, MA), and TRPV4 (1:400; Waltham, MA) were diluted 1:1 (blocking buffer: PBS) and applied overnight at 4°C. The slides and coverslips were washed 3x in PBS (5 min each) and then incubated for 1 h at RT (in the dark) with Alexa Fluor 488- and 594-conjugated secondary antibodies (Invitrogen; 1:1000 dilution). Nuclei were counterstained with DAPI (1:3000), and samples were mounted with FluorSave™ mounting media (Sigma Aldrich, St. Louis, MO). Retinal sections and cell coverslips were imaged via a Nikon Ti-Eclipse fluorescence microscope (Nikon Instruments, Melville, NY) with a 20x objective. Both central and peripheral retina regions were captured for quantitative analysis.

### Image analysis

The fluorescence images were analyzed via ImageJ (v1.53c; NIH). Regions of interest (ROIs) were defined based on DAPI staining. Binary masks of each ROI were generated, and the mean fluorescence intensity was measured in the corresponding channels. Negative control sections were used to define the background signal which was subtracted from all measurements.

### Liquid Chromatography**lJ**Tandem Mass Spectrometry

Primary and immortalized Müller glia were collected following 24 h of incubation with or without the TRPV4 agonist. Müller glial cultures and conditioned media were collected in LC-MS-grade methanol for analysis and quantification of the eicosanoid and PUFA pathways via LC_MS/MS via our previously published protocol [33]. A total of 400 pg of each deuterated internal standard (prostaglandin E_2_-d_4_, 15-HETE-d_8_, leukotriene B_4_-d_4_, AA-d_8_, docosahexaenoic acid-d_5_, or LXA_4_-d_5_) was added to each sample to calculate the extraction recovery of each lipid class. The supernatants containing lipids were extracted via C18 solid-phase columns and analyzed via a triple-quadrupole linear ion-trap LC_MS/MS system (AB SCIEX 4500 QTRAP) equipped with a Kinetex C18 mini-bore column. The mobile phase was a linear gradient of A (water, acetonitrile, and acetic acid (72:28:0.01 by volume)) and B (isopropanol and acetonitrile (60:40 by volume)), with a flow rate of 450 µL/min. MS/MS analysis was performed in negative ion mode. PUFAs and lipoxygenase pathway metabolites were identified and quantified via scheduled multiple reaction monitoring (MRM) using 4–6 specific transitions for each analyte. Established diagnostic fragment ions were used for quantification: arachidonic acid (AA; 303→259 m/z), 5-hydroxyeicosatetraenoic acid (5-HETE; 319→115 m/z), 15-HETE (319→175 m/z), 12-HETE (319→179 m/z), LXA_4_ (351→115 m/z), LXB_4_ (351→221 m/z), and DHA (327→283 m/z) EPA (301→257). Peaks were identified based on the integration criterion of a signal-to-noise ratio of at least 5:1. Class-specific deuterated internal standards were used to validate the retention times of the analytes in each sample. Quantification was based on analyte-specific calibration curves prepared from synthetic standards (Cayman Chemical, Ann Arbor, MI) [34]. The results were normalized to the number of cells (1 × 10^6^ cells/sample).

### Flow Cytometry

Primary Müller glia were treated with a TRPV4 agonist and/or LXB_4_ for 24 h and then harvested with TrypLE Express (Gibco, Waltham, MA). The cells were washed in ice-cold PBS and fixed/permeabilized. Fixed cells were incubated overnight at 4°C. The cells were washed in wash buffer and then stained with primary antibody for 1 h at RT in the dark, followed by washing with cell staining wash buffer. The cells were incubated with secondary antibody for 1 h at RT, washed, and resuspended in wash buffer for analysis via flow cytometry. The cells were analyzed via an Attune Acoustic Focusing Flow Cytometer (Thermo Fisher, Waltham, MA). Dead cells and debris were excluded by forward/side scatter gating. Singlets were gated by FSC-A vs. FSC-H. GFAP⁺ cells were identified by comparison to isotype controls. The data were analyzed with FlowJo v10 (BD, Ashland, OR). The fraction of GFAP-positive cells was used for quantitative comparisons.

### Phospho-kinase Assay

For phospho-kinase profiling, the Proteome Profiler Human Phospho-Kinase Array (R&D Systems, Minneapolis, MN) was used. rMC-1 cells were treated with or without 1 µM LXB_4_ for 30 min and then lysed in the kit’s Lysis Buffer 6 supplemented with protease inhibitors, according to the manufacturer’s instructions. Protein concentrations were determined using a BCA assay (Pierce, Thermo Fisher), and 200 µg of total protein per sample was diluted to 2 mL in Array Buffer 1. Membranes were blocked in Array Buffer 1 for 1 h at RT and incubated with diluted lysates overnight at 2–8 °C with rocking. After washing 3x, membranes were incubated with the biotinylated detection antibody cocktail (Parts A and B) for 2 h at RT, washed 3x, and incubated with IRDye 800CW Streptavidin (LI-COR Biosciences, Lincoln, NE) for 30 min. Membranes were washed 3x, and then imaged on a LI-COR Odyssey system. Spot intensities were quantified using LI-COR software after background subtraction and were normalized to the array’s positive control signals.

### LXB_4_ treatment

For single-cell transcriptomic analysis, the mice were treated *i.p.* with LXB_4_ methyl ester (Cayman Chemical, Ann Arbor, MI) (1 µg in 100 µL of PBS) once daily and topically (1 µg in 5 µL of PBS per eye) 3x daily for 3 days. Vehicle (ethanol) for LXB_4_ methyl ester was removed under a stream of nitrogen, and LXB_4_ methyl ester was resuspended in sterile PBS immediately prior to injection or topical treatment. On day 4, the mice were euthanized for retinal dissociation and single-cell library preparation. For IHC analysis, the mice received daily LXB_4_ methyl ester (1 µg in 100 µL of PBS) via *i.p.* injection and topical ocular administration for 1 wk. The control mice received equivalent volumes of sterile PBS. LXB_4_ was administered 15 min prior to OHT induction.

### Single-cell transcriptomics

Single-cell transcriptomic data were reanalyzed for the Müller glial population from a previously published dataset from our laboratory[24]. Retinas were dissociated into single-cell suspensions via a papain dissociation kit (Worthington, Columbus, OH), followed by rod photoreceptor depletion in a magnetic column via CD133-biotin labeling and anti-biotin magnetic beads (Miltenyi Biotech, Bergisch Gladbach, Germany). Single cells were loaded on the 10x Chromium Single Cell 3′ v3.1 platform (10x Genomics, Pleasanton, CA) to generate barcoded gel-bead emulsions, and captured RNA was reverse-transcribed and amplified into cDNA libraries, which were quality-checked on an Agilent Bioanalyzer. Libraries were sequenced on an Illumina NovaSeq S1 100SR flow cell, and raw BCL files were demultiplexed via Illumina bcl2fastq2 at the QB3 Genomics Core, UC Berkeley, Berkeley, CA (RRID: SCR_022170). The sequencing reads were aligned to the mm10 (mouse) genome with Cell Ranger software (10x Genomics, Pleasanton, CA). Downstream analyses were performed in Seurat (v4.1.0) as described previously [24,35]. Briefly, following quality control on the basis of the percentage of mitochondrial transcripts, the data were normalized via the normalizeData() function. Variable features were identified with FindVariableFeatures() and integrated across datasets via FindIntegrationAnchors() and IntegrateData() on the first 30 principal components (PCs). Postintegration, the data were scaled with ScaleData(), clustered with FindNeighbors() and FindClusters() (resolution = 0.8) on the first 30 PCs and dimensionally reduced via RunUMAP() on the first 30 PCs. Retinal cell types were annotated by identifying cluster-specific markers with FindAllMarkers() and comparing them to known markers [35]. Differential gene expression in Müller glia was determined via FindMarkers() and visualized with VlnPlot() and DotPlot().

### Western blot

Treated Müller glia were lysed in RIPA buffer (Thermo Fisher, Waltham, MA) containing a protease and phosphatase inhibitor cocktail (Thermo Fisher, Waltham, MA) at a 1:100 dilution. Protein concentrations were determined via a BCA assay (Pierce, Thermo Fisher). Equal amounts of protein (20 µg) were denatured, separated on 10% SDS_PAGE gels, and then transferred to nitrocellulose membranes (Bio-Rad Laboratories, Hercules, CA). The membranes were blocked in Intercept Blocking Buffer (LiCOR, NE) for 1 h at RT and then probed overnight at 4°C with the following antibodies: anti-phospho-STAT3 (Ser727; rabbit monoclonal; 1:1000; Abcam, Cambridge, MA), total anti-STAT3 (rabbit monoclonal EPR361; 1:2000; Abcam, Cambridge, MA), and anti-GAPDH (mouse monoclonal; 1:10,000; Cell Signaling Technology, Dancars, MA). After 3 washes in TBST, the membranes were incubated with IRDye 800CW donkey anti-rabbit IgG and IRDye 680RD donkey anti-mouse IgG (1:10,000; LI-COR Biosciences, Lincoln, NE) for 1 h at RT in the dark. The blots were imaged on a LI-COR Odyssey system. Band intensities were quantified with ImageJ (NIH). Phospho-STAT3 signals were normalized to total STAT3 and GAPDH signals and normalized to those of untreated controls for fold-change calculations.

### Statistical analysis

All the data are presented as the means ± standard errors of the means (SEMs). Biological replicates (n) represent the number of animals (for *in vivo* experiments) or independent cell culture experiments (for *in vitro* assays). Statistical analyses were conducted in GraphPad Prism 9.0 (GraphPad Software, San Diego, CA). For two-group comparisons, an unpaired Student’s t-test was used. For multiple-group comparisons, one-way analysis of variance (ANOVA) followed by Tukey’s post hoc test was performed, with the assumption of a normal distribution of variance. Statistical significance was defined as p < 0.05.

## Results

### Müller glia express a functional lipoxin biosynthetic pathway

Our previous studies established that astrocytes generate lipoxins and that LXB_4_ treatment reduces Müller glia reactivity [23,36]. We next asked whether Müller glia themselves synthesize lipoxins. Primary mouse Müller glia were isolated and validated by ICC for canonical markers, including vimentin, glutamine synthetase, and SOX9, and were confirmed to be negative for GFAP and IBA1 (Fig. 1A). Immortalized rMC-1 cells were validated via the same markers and expressed all the same positive and negative markers (Supplementary Fig. 1A). Bulk RNA-seq analyses of cultured Müller glia revealed robust expression of *Alox5*, along with significant levels of *Alox5ap, Alox12* and *Alox1*5 (Fig. 1B), indicating the presence of key lipoxygenase pathway enzymes for lipoxin generation. Targeted lipidomic profiling revealed the presence of arachidonic acid (AA) and its lipoxygenase metabolites 5-HETE, 12-HETE, and 15-HETE in both cell lysates and conditioned media from both primary (Fig. 1C) and rMC-1 cells (Supplementary Fig. 1B). qPCR analysis established robust expression of the *Alox5* (Ct: 27.18) and *Alox15* (Ct: 28.05) genes in primary Müller glia, confirming the RNA-seq data (Fig. 1D). More importantly, Müller glia generated significant amounts of endogenous LXB_4_ (114 pg/mL) and its pathway markers 5-HETE (1210 ng/mL) and 15-HETE (1502 ng/mL) (Fig. 1E). Together, these findings establish that primary Müller glia functionally express the LXB_4_ biosynthetic pathway (*Alox15* and *Alox5)*, revealing a previously unrecognized glial lipid mediator pathway.

**Figure 1.**
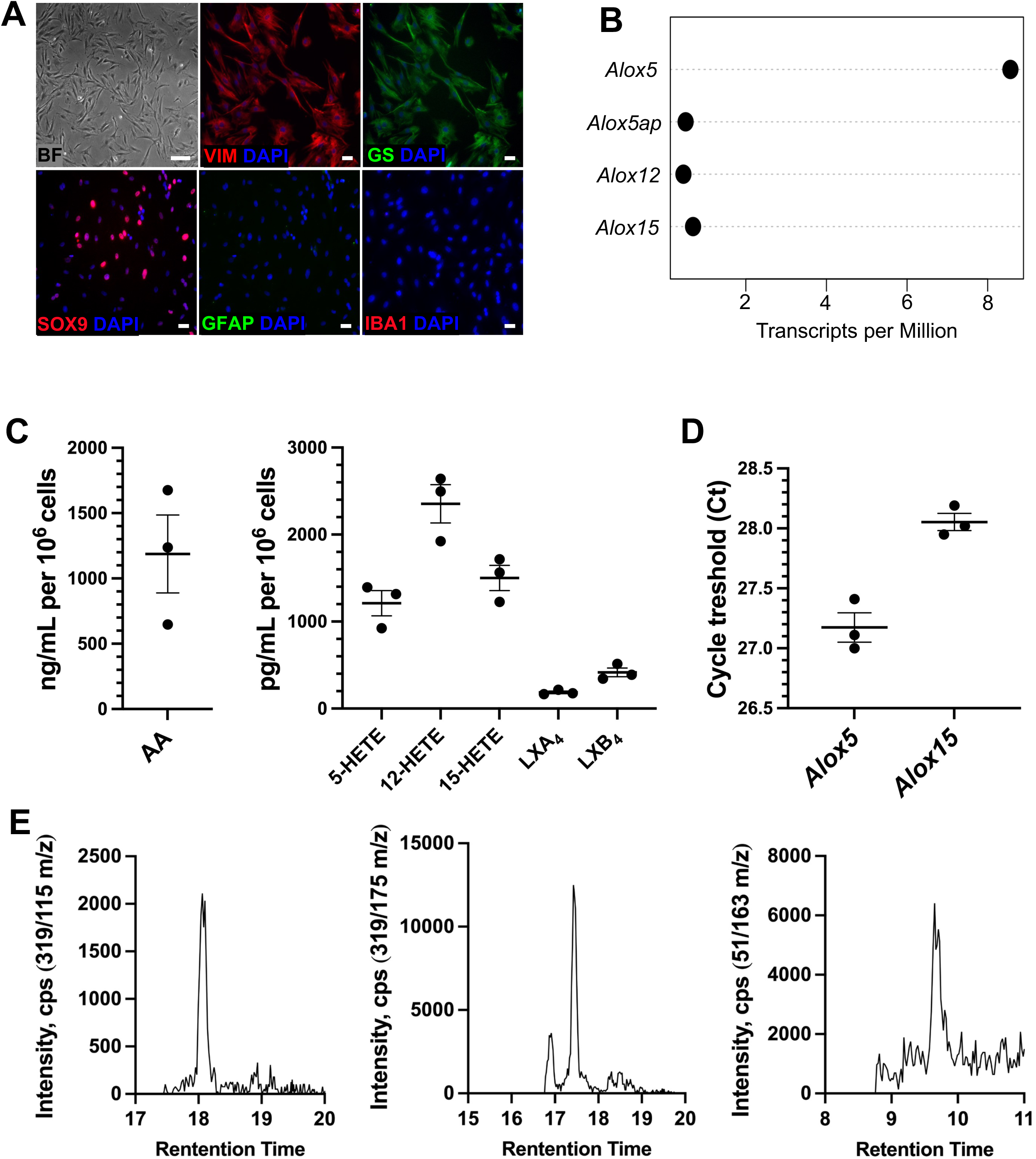
Functional expression of the lipoxin pathway in Müller glia. **(A)** Primary mouse Müller glia were stained for canonical markers (vimentin [VIM], glutamine synthetase [GS], and SOX9) and negative control markers (GFAP and IBA1). **(B)** Bulk RNA sequencing of primary Müller glia for lipoxygenase pathway genes (n = 3). **(C)** LC_MS/MS quantification of released eicosanoid substrate (AA) and corresponding lipoxygenase products in primary Müller glia (n = 3). **(D)** qPCR analysis of *Alox5* and *Alox15* expression in primary Müller glia (n = 3). **(E)** Representative diagnostic raw chromatograms and LC retention times for Müller glia-generated 5-HETE, 15-HETE, and LXB_4_.

### Ocular hypertension induces a reactive Müller glial phenotype and increases TRPV4 expression

To model sustained mechanical insult, we induced chronic OHT in C57BL/6J mice via anterior-chamber silicone-oil injection [23,24,28]. Mice were euthanized, and the retinas were collected at 2, 6, and 8 wk post-injection to assess Müller glia reactivity by IHC with GFAP as an established marker (Fig. 2A). Imaging revealed an 80% increase in GFAP signal intensity in the ganglion cell layer (GCL) and inner nuclear layer (INL), specifically in astrocytes and Müller glia, at 2 wk and 6 wk (Fig. 2B). In parallel, the TRPV4 signal intensity increased significantly at every time point examined (Fig. 2C), with a 114.6% increase in TRPV4 expression relative to that in age-matched normotensive controls at 6 wk (Fig. 2D). qPCR analysis at 4 wk OHT revealed the upregulation of *Il-6* (6.2-fold), *Tnf-*α (20-fold), and *Stat3* (25.5-fold) when compared with controls (Fig. 2E), which was consistent with proinflammatory activation. Gene set enrichment analysis (GSEA) of RNA-seq data from MACS-isolated Müller glia supported these findings, revealing significant upregulation of cytokine-related pathways, including *Il-6* and *Tnf-*α production. In addition, pathways related to visual perception and the response of sensory stimuli to light stimuli were also downregulated for the primary supportive cell of neuronal cells, indicating possible negative changes in Müller glia supportive functional pathways (Fig. 2F). These data demonstrate that chronic OHT induces reactive gliosis, TRPV4 upregulation, and inflammation, highlighting the role of Müller glia as central responders to mechanical insult.

**Figure 2.**
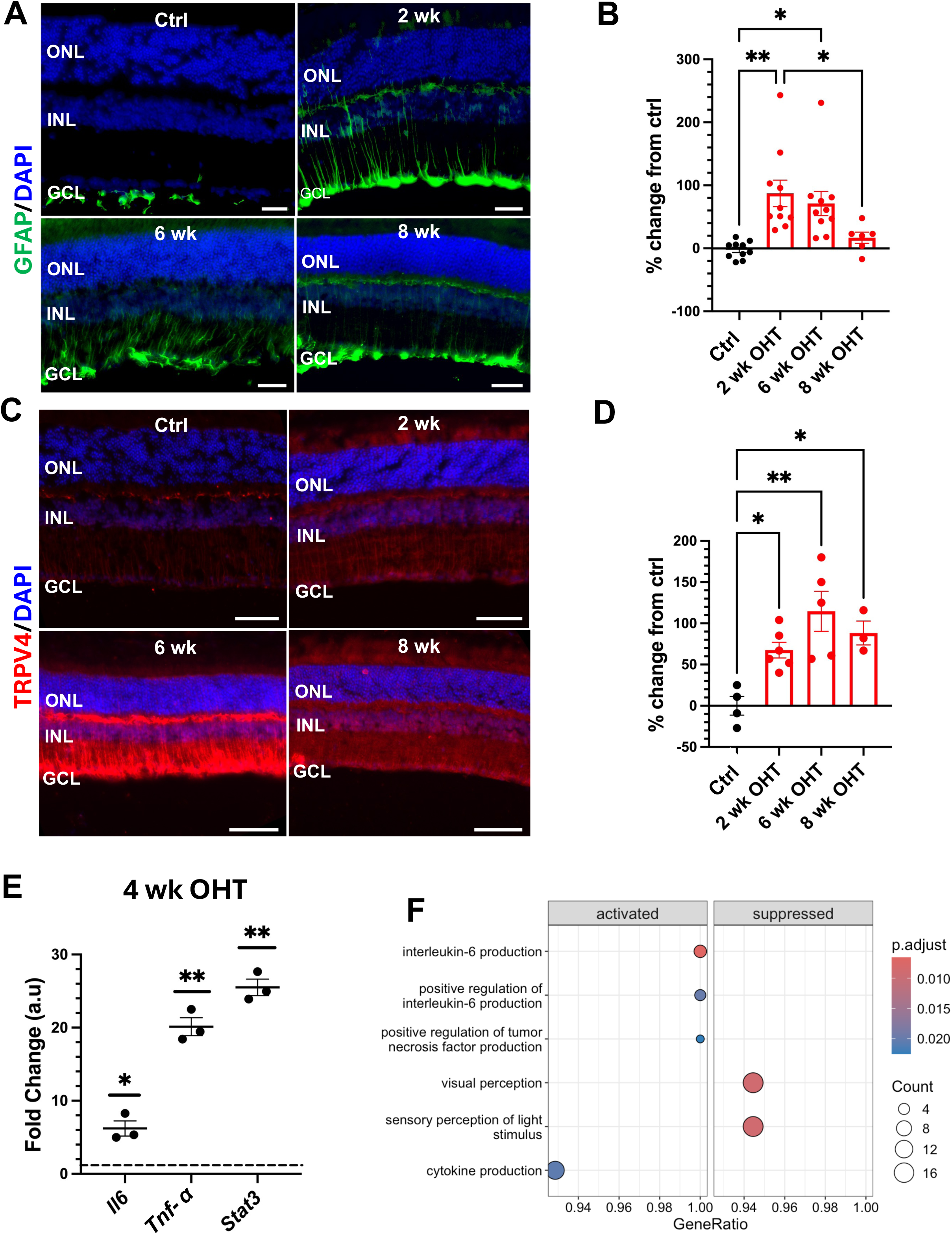
Ocular hypertension (OHT) induces GFAP and TRPV4 expression, which is correlated with a proinflammatory phenotype. **(A)** Representative IHC images of retinal sections from mice with sustained OHT at 2, 6, and 8 wk post-injection and age-matched normotensive controls stained for GFAP and DAPI. **(B)** Quantification of GFAP immunoreactivity, expressed as the percent change from the control (n = 5-10). **(C)** Representative immunofluorescence images from mice with (OHT) and normotensive controls stained for TRPV4 and DAPI. **(D)** Quantification of TRPV4 immunoreactivity, expressed as the percent change from control (n = 3–7). **(E)** qPCR analysis of *Il6, Tnf-*α, and *Stat3* in whole retinas collected 4 wk post-OHT (n = 3). Data were normalized to *Gapdh* and matched with normotensive controls. **(F)** Gene set enrichment analysis (GSEA) of MACS-isolated Müller glia from chronic OHT conditions. Images acquired at 20x magnification; scale bar = 50 µm. GCL = ganglion cell layer, INL = inner nuclear layer, ONL = outer nuclear layer. Statistical analysis: one-way ANOVA with Tukey’s post hoc test (*p<0.05, **p<0.01, ***p<0.001) or unpaired Student’s t-test (*p<0.05, **p<0.01).

### TRPV4 activation drives Müller glia reactivity and amplifies LXB_4_ production

To isolate the specific contribution of TRPV4 signaling, primary and immortalized Müller glia were treated for 24 h with the selective TRPV4 agonist GSK. Cells from control (ctrl) and treated conditions were analyzed by flow cytometry, and GFAP^+^ cells were selectively gated (Fig. 3A). Quantification revealed that TRPV4 activation caused a 118% increase in GFAP-positive cells compared with vehicle-treated controls (DMSO) (Fig. 3B). IHC further confirmed the increased expression of GFAP and vimentin along Müller glial filaments following TRPV4 activation (Fig. 3C). Lipidomic analyses revealed that TRPV4 activation triggered a marked increase in the release of the lipoxygenase substrate arachidonic acid (AA) and 12-HETE a downstream signature 12/15-LOX metabolite. Notably, reactive Müller glia increased LXB_4_ production by 2.3-fold relative to controls (Fig. 3D). Thus, TRPV4 activation is sufficient to drive Müller glial reactivity and activate lipoxygenase pathways to amplify intrinsic LXB_4_ production.

**Figure 3.**
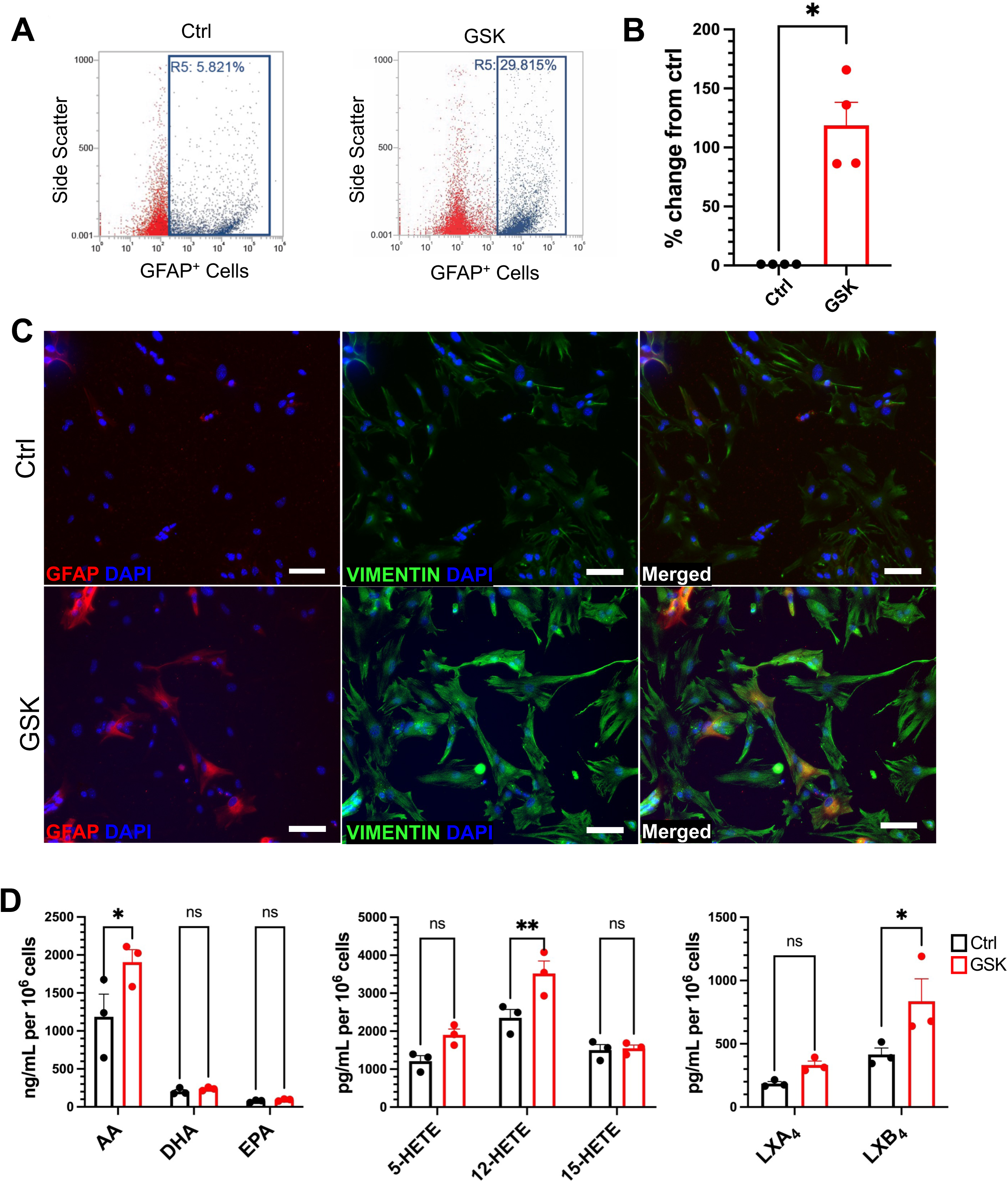
Activation of TRPV4 induces Müller glial reactivity and enhances LXB_4_ formation. **(A-B)** Flow cytometry gating and quantification of GFAP⁺ cells following TRPV4 agonist (GSK) vs vehicle (DMSO, Ctrl) treatment (n = 4). **(C)** Representative ICC images of GFAP and vimentin in GSK- or vehicle-treated (Ctrl) Müller glia. **(D)** LC_MS/MS quantification of released PUFAs, 5-LOX, 12/15-LOX metabolites, and lipoxins (LXA_4_, LXB_4_) by Müller glia following GSK treatment (n = 3). Images acquired at 20x magnification; scale bar = 50 µm. Statistical analysis: unpaired Student’s t test (*p<0.05, **p<0.01, ***p<0.001).

### LXB_4_ suppresses JAK-STAT signaling in homeostatic Müller glia

Because IL-6/JAK/STAT is a central driver of reactive gliosis and was upregulated in our OHT model (Fig. 2E), we next investigated whether exogenous LXB_4_ modulates this pathway. When immortalized Müller glia (rMC-1) were treated with LXB_4,_ a phosphokinase array revealed that LXB_4_ selectively reduced the phosphorylation of STAT2-Y690 (18%), STAT5a/b-Y699 (16.8%), and STAT3-S727 (19%) (Fig. 4A). To assess the transcriptional consequences of LXB_4_ signaling *in vivo*, retinas from LXB_4_ treated mice were analyzed by single-cell RNA-seq. UMAP analysis revealed a distinct Müller–glial cluster (Fig. 4B), within which *Stat1* and *Stat3* transcript levels were reduced in LXB_4_-treated mice (Fig. 4C). Differential gene expression analysis from single-seq data further revealed downregulation of *Jak2*, *Il6st*, *Stat5b*, and *Stat2* (Fig. 4D), indicating broad suppression of the inflammatory JAK-STAT network. Collectively, these findings demonstrate that LXB_4_ broadly attenuates JAK-STAT signaling at both the protein and transcriptional levels in homeostatic Müller glia.

**Figure 4.**
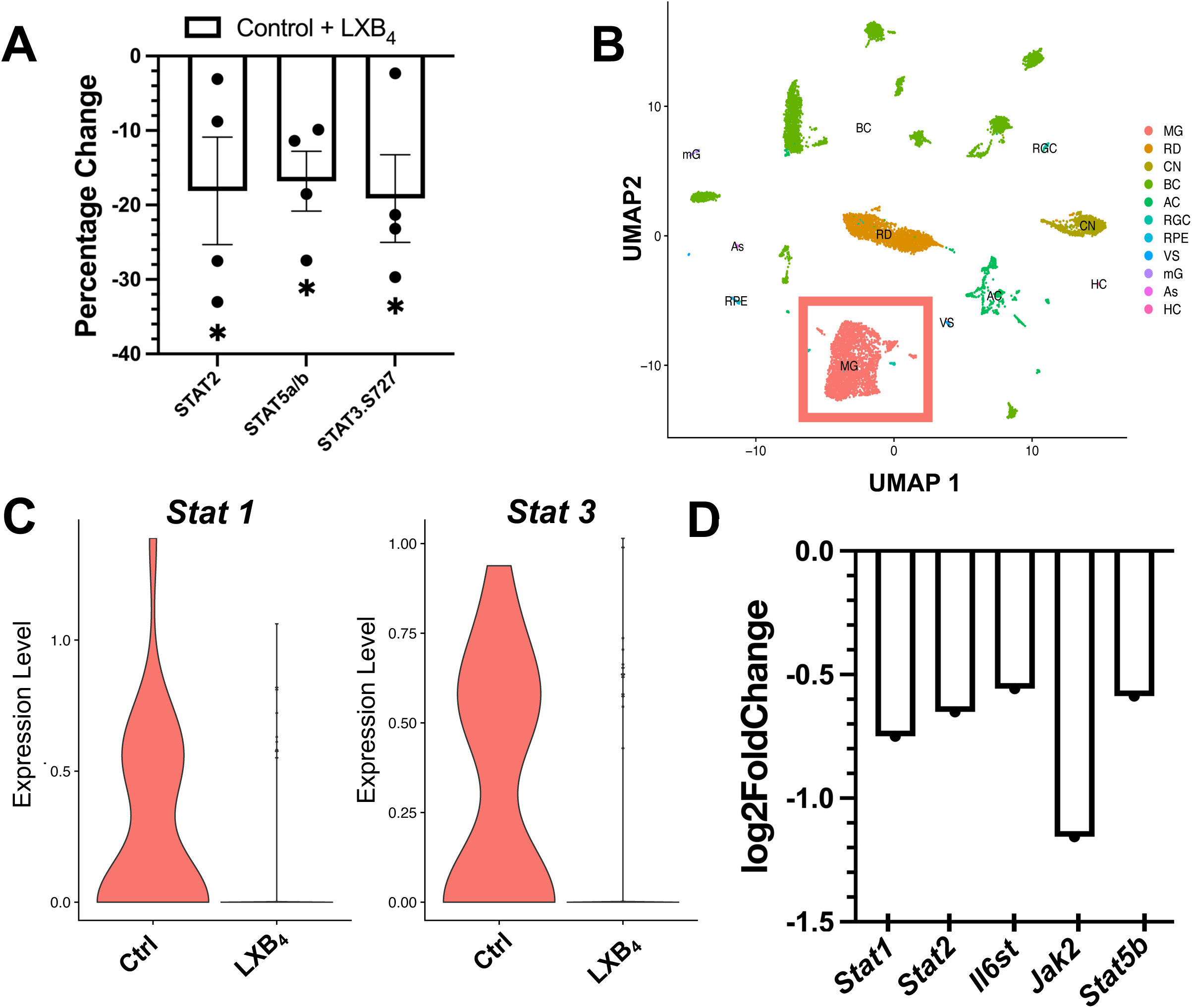
LXB_4_ Regulates JAK-STATs in Homeostatic Müller Glia *In Vitro* and *In Vivo*. **(A)** Phospho-kinase array of immortalized Müller glia (rMC-1) treated with or without LXB_4_ (n = 4). **(B)** UMAP projection of single-cell RNA sequencing data from retinas treated with or without LXB_4_, highlighting the Müller glia (MG) populations. **(C)** Violin plots comparing Stat1 and Stat3 expression in MG populations between groups**. (D)** Differential gene expression of *Jak2, Stat2, Stat5b, Il6st,* and *Stat1* in MGs from the scRNA-seq dataset. Statistical analysis: unpaired Student’s t test (*p<0.05).

### LXB_4_ inhibits TRPV4-induced Müller glial reactivity via IL-6/STAT3 suppression

To determine whether LXB_4_ can prevent TRPV4-induced Müller glial reactivity, we pretreated primary Müller glia with LXB_4_ prior to stimulation with the TRPV4 agonist GSK. After 24 h, flow cytometry analysis revealed that LXB_4_ pretreatment significantly reduced the number of GFAP-positive cells by 84.6% compared with that in the GSK-treated controls (Fig. 5A-B). ICC further confirmed that the GFAP and vimentin filament intensities significantly decreased, returning to near-baseline levels (Fig. 5C). qPCR analysis revealed a significant decrease in *Il6* expression in LXB_4_-treated primary Müller glia (Fig. 5D). This reduction in *Il6* (1.6-fold) was accompanied by decreased STAT3 phosphorylation (Ser727) by 75%, as determined by Western blotting and normalized to total STAT3 and GAPDH (Fig. 5D), which is consistent with *Il6* acting upstream of STAT3 activation. Together, these results demonstrate that LXB_4_ effectively suppresses TRPV4-induced inflammatory activation in immortalized Müller glia by downregulating *Il6* expression, inhibiting STAT3 phosphorylation, and preventing the upregulation of cytoskeletal reactivity markers.

**Figure 5.**
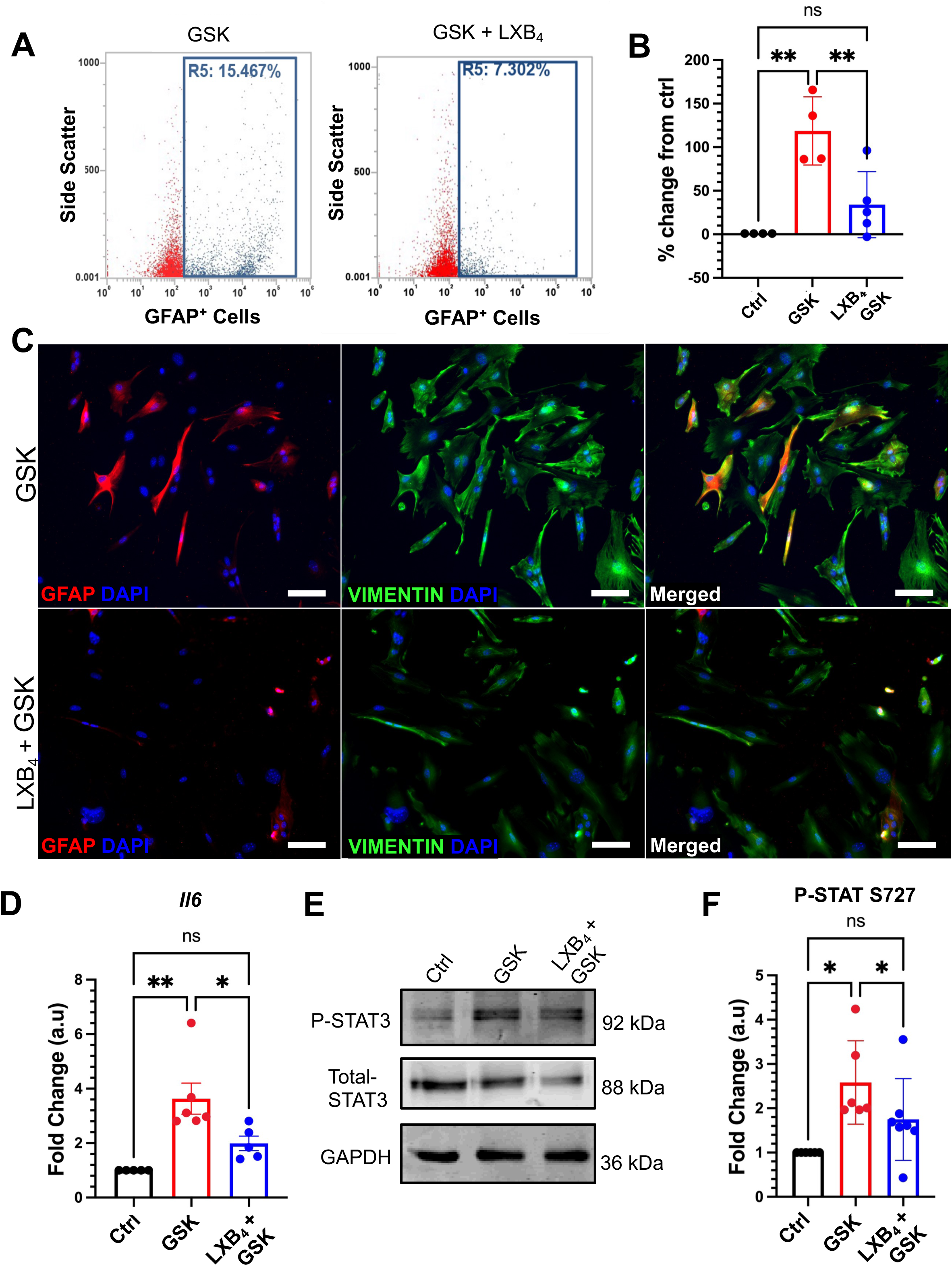
LXB_4_ Suppresses the IL6-STAT Pathway *In Vitro*. **(A-B)** Flow cytometry of GFAP⁺ cells in control (vehicle), GSK, or GSK + LXB_4_ treated primary Müller glia (n = 4). **(C)** ICC images of GFAP and vimentin in the GSK, or GSK+LXB_4_ treated cells. **(D)** qPCR results for *Il6* expression in control, GSK, or LXB_4_ + GSK treated Müller glia (n = 5). **(E-F)** Western blot analysis of phospho-STAT3 (S727) and total STAT3. Representative blots are shown; GAPDH was used as a loading control (n = 5). Images acquired at 20x magnification; scale bar = 50 µm. Statistical analysis: one-way ANOVA with Tukey’s post hoc test (*p<0.05, **p<0.01).

### LXB_4_ inhibits inflammatory gene expression and TRPV4 upregulation in chronic and acute OHT models

Finally, we evaluated the therapeutic potential of LXB_4_ in both chronic and acute models of OHT [23,24]. In the chronic model, qPCR analysis of whole retina RNA from mice treated with LXB_4_ after 4 wk of OHT revealed marked reductions in *Stat3* (13.2-fold), *Il6* (4.5-fold), and *TNF-*α (14.7-fold) expression when directly compared to vehicle-treated mice with OHT (Fig. 6A). In a separate acute OHT model (1 wk post-induction), IHC demonstrated that LXB_4_ treatment prevented the OHT-induced increase in TRPV4 expression, particularly in the GCL and INL (Fig. 6B). TRPV4 signal intensity was reduced by 63% in LXB_4_-treated mice when directly compared to vehicle-treated mice with OHT (Fig. 6C). These *in vivo* results reinforce the therapeutic neuroprotective potential of LXB_4_, as it suppresses inflammation and TRPV4 upregulation in both acute and chronic OHT models of glaucomatous neurodegeneration.

**Figure 6.**
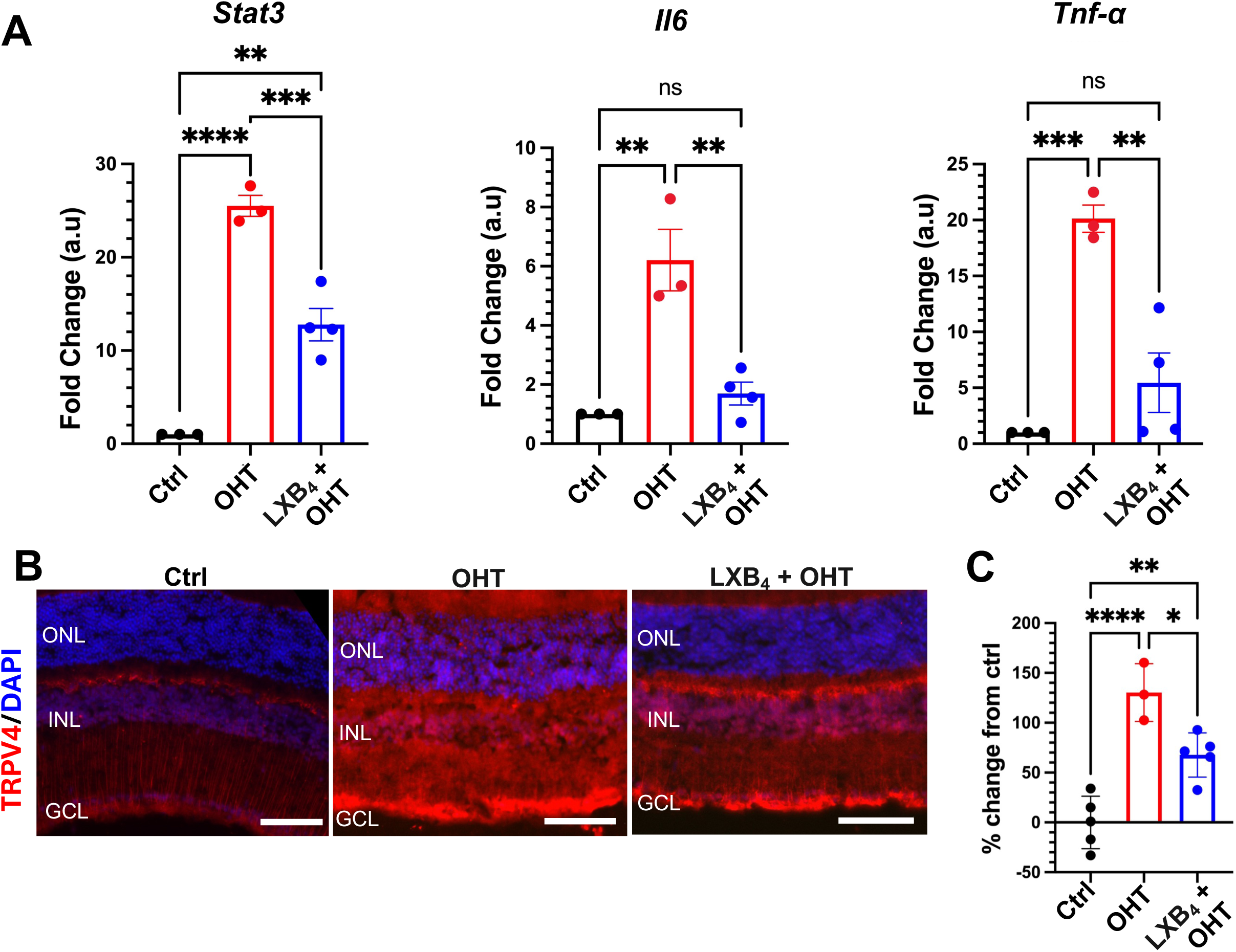
LXB_4_ Treatment Suppresses Proinflammatory Markers and TRPV4 Expression *In Vivo*. **(A)** qPCR analyses of *Stat3, Il6*, and *TNF-*α expression in whole retinas of normotensive eyes and at 4 wk of chronic OHT with or without LXB_4_ treatment (n = 4). **(B)** Representative IHC images of TRPV4 IHC in normotensive controls and those subjected to acute OHT for 1 wk with or without LXB_4_ treatment. **(C)** Quantification of TRPV4 protein expression in control retinas and OHT-treated retinas with or without LXB_4_ treatment. Protein levels were quantified via Western blotting, with GAPDH used as a loading control (n = 3–5). Images acquired at 20x magnification; scale bar = 50 µm. Statistical analysis: one-way ANOVA with Tukey’s post hoc test (*p<0.05, **p<0.01, ***p<0.001, ****p<0.0001).

## Discussion

The role of Müller glia in glaucoma pathogenesis is complex and multifaceted, transitioning from an initially protective response to a chronic, maladaptive phenotype that accelerates RGC loss [37]. Building on prior work showing that astrocyte-derived LXB_4_ protects RGCs, we extended this paradigm to Müller glia [23,38]. We show that these cells are both sensors of biomechanical insult via TRPV4 and active producers of an intrinsic, lipoxin-based anti-inflammatory circuit. These findings suggest that Müller glia not only contribute to neuroinflammation but also possess a self-regulatory mechanism that could be amplified to suppress gliosis and protect RGCs.

We discovered that Müller glia express a functional lipoxin biosynthetic circuit characterized by 5-LOX and 15-LOX expression and the endogenous production of LXB_4_. While previous studies have identified retinal astrocytes as a source of lipoxins [23], our findings extend this paradigm by demonstrating that Müller glia, the most abundant macroglia cell type in the retina, also is a key contributor to this neuroprotective retinal pathway. LC_MS/MS-based lipidomics confirmed that these cells not only express the necessary enzymes but also actively produce key lipoxin precursors (5-, 12-, and 15-HETE) and release neuroprotective LXB_4_ under homeostatic conditions, revealing a previously unrecognized biosynthetic capacity.

*In vivo*, chronic OHT robustly upregulated TRPV4 in Müller glia, leading to the activation of canonical gliosis hallmarks, including IL-6/STAT3 signaling and GFAP expression. These findings align with those of previous studies in other central nervous system (CNS) models, which demonstrated that TRPV4 activation leads to Ca²⁺-dependent p-STAT3 induction and that inhibiting STAT3 mitigates neurodegeneration [39–41]. Our *in vitro* data further support this link, showing that pharmacological TRPV4 activation in Müller glia recapitulates gliotic changes while increasing lipoxin production. These findings suggest a direct functional relationship between the biomechanical stress sensor TRPV4 and the gliotic response, highlighting a key regulatory point for Müller glia function.

TRPV4-activated Müller glia reactivity induced both inflammatory (IL-6 and TNF-α) and neuroprotective (LXB_4_) pathways. The lipoxygenase pathways (5-LOX and 12/15-LOX) that were amplified, in addition to generating protective lipoxins, can also generate pro-inflammatory lipid mediators [16,42]. 5-LOX, a rate limiting enzyme for lipoxin generation, can also generate pro-inflammatory leukotrienes. However, *in vitro* primary and immortalized Müller glia did not generate any detectable amounts of leukotrienes under any conditions. Consistent with our previous lipidomic analyzes of healthy and glaucomatous retinas [23,38], these findings are consistent with our previous *in vivo* studies that detected no leukotrienes in the retina or optic nerve in healthy eyes or eyes with OHT (JCI, Shruthi ref). 12-HETE, which is a primary product of the mouse 12/15-LOX (Alox15) but of the human 15-LOX (ALOX15) was increased in reactive rodent Müller glia [43]. 12-HETE has been reported in patients with diabetic retinopathy and is known to activate and upregulate inflammatory markers, including IL-6 and TNF-α, in Müller glia [43,44]. These findings indicate that TRPV4-induced gliosis triggers lipoxygenase pathways to amplify neuroprotective signaling (i.e., LXB_4_) but could also initiate an inflammatory pathway (i.e., 12-HETE). Hence, TRPV4 activation triggered by OHT-induced biomechanical stress triggers gliosis but also can increase lipoxin biosynthesis in Müller glia, which indicates a self-regulating mechanism where TRPV4-driven gliosis is counterbalanced by amplified endogenous lipoxin signaling.

LXB_4_ treatment disrupted the inflammatory circuit and restored homeostatic balance under chronic OHT. Our experiments revealed that LXB_4_ treatment suppresses STAT3 phosphorylation, downregulates TRPV4 expression, and reverses gliosis (Figs. 4–6), effects that parallel those observed with pharmacological TRPV4 inhibition [9]. Our findings align with studies that have shown treatment with omega-3 pro-resolving lipid mediators, such as Maresin-1, has overlapping anti-inflammatory actions by decreasing TRPV4 expression in other CNS models [45]. Lipoxins, unlike Maresins, are an endogenous protective lipid mediator pathway in the retina. Hence, therapeutic amplification of LXB_4_ signaling selectively targets the resident lipid mediator pathway in the retina. Stable analogs of lipoxin have been developed and especially the protective action of LXA_4_ mimetics are well characterized in multiple models of CNS dysfunction [46]. In models of posterior uveitis, a form of retinal inflammation, both LXA_4_ and LXB_4_ reduce macroglial reactivity, microglial activation, and macrophage infiltration [47]. However, LXB_4_’s neuroprotective actions in OHT and neurotoxic models of glaucomatous injury are superior those of LXA_4_ (JCI Ref). While prior studies have focused on lipoxin activity in modulating astrocyte and microglial inflammatory responses, our current findings are the first to identify Müller glia as both a source and a target of lipoxin signaling, establishing a unique self-regulatory inflammatory circuit specific to the retina [23,24,36,38].

The identification of a counter-regulatory lipoxin circuit in Müller glia has potential translational implications for glaucoma treatment. Since Müller glia are the major glial cell type in the retina and originate from the same progenitor cells as RGCs do, they represent an ideal target for therapeutic intervention [7,48]. Therefore, strategies that augment this intrinsic protective circuit could be a targeted approach to prevent the maladaptive gliosis that precedes RGC degeneration. For example, this could be achieved pharmacologically with established stable LXB_4_ analogs or genetically via macroglia specific AAV-mediated amplification of the lipoxin pathway. This targeted approach is particularly promising, as it leverages the retina’s own neuroprotective mechanisms to restore homeostatic balance.

This study has several limitations. While our findings provide mechanistic insight into Müller glia reactivity, longer-term studies are needed to evaluate the impact of LXB_4_ treatment on the homeostatic function of Müller glia. Although we focused on TRPV4 and STAT3 signaling, other inflammatory pathways, such as the NF-κB or MAPK pathways, likely contribute to the gliotic response and deserve further exploration [49,50]. Additionally, much of our mechanistic work was performed in primary and immortalized Müller glia cultures. While these systems are invaluable for dissecting signaling pathways in a controlled environment, they inevitably simplify the cellular complexity of the retina. Important interactions between glia, neurons, and the vasculature are not fully understood *in vitro*. Future work using conditional knockout models and live imaging will help clarify the temporal dynamics of this lipid signaling axis *in vivo*.

In summary, we identified the intrinsic neuroprotective and anti-inflammatory lipoxin circuit in Müller glia, which are the primary and most abundant supportive glia in the retina. We established that chronic activation of the mechanosensitive TRPV4 ion channel in Müller glia drives sustained gliosis and IL-6/STAT3 pathway activation. This TRPV4-induced gliosis is counter-regulated by the amplification of Müller glia LXB_4_ formation. LXB_4_ targets both OHT-induced TRPV4 overexpression and TRPV4-induced STAT3 phosphorylation, thereby reversing the inflammatory reactivity of Müller glia. Hence, targeting the Müller glia lipoxin pathway could provide new therapeutic opportunities for glaucoma and other retinal neurodegenerative diseases associated with chronic activation of TRPV4 and gliosis of Müller cells.

## Declarations

### Ethics approval and consent to participate

All animals were handled in accordance with the ARVO Statement for the Use of Animals in Ophthalmic and Vision Research, and all procedures were approved by the Institutional Animal Care and Use Committee (IACUC) at the University of California, Berkeley.

### Consent for publication

Not applicable

### Availability of data and materials

The MACS sorted Müller glia and primary Müller glia datasets generated in this study was deposited in the GEO (GSE306785, RNAseq). The single cell dataset generated in this study was deposited in the GEO (GSE251716, scRNA-seq.)

## Funding

This work was partly supported by the NIH training grants 5T32EY007043-45, R01EY030218, and P30EY003176.

### Authors and contributions

M.K., J.G.F., and K.G. conceived the study; M.K., S.K., S.M., and R.N. performed the experiments. M.K. and R.N. performed the data analysis; M.K., S.K., J.G.F., and K.G. wrote the manuscript.

## Acknowledgments

We acknowledge the QB3 Genomics core RRID: SCR_022170, UC Berkeley, for sequencing support, and the CRL Molecular Imaging Center, RRID: SCR_017852, UC Berkeley, for imaging and image analysis support.

## Abbreviations

AA: Arachidonic acid
CNS: Central Nervous System
DHA: Docosahexaenoic acid
GCL: Ganglion cell layer
GFAP: Glial fibrillary acidic protein
GSEA: Gene set enrichment analysis
HETE: Hydroxyeicosatetraenoic acid
ICC: Immunocytochemistry
IHC: Immunohistochemistry
IL-6: Interleukin 6
INL: Inner nuclear layer
IOP: Intraocular pressure JAK Janus Kinases
LC–MS/MS: Liquid chromatography–tandem mass spectrometry
LXB_4_: Lipoxin B_4_
OHT: Ocular hypertension
ONL: Outer nuclear layer
PUFA: Polyunsaturated fatty acid
RGC: Retinal ganglion cell
STAT: Signal transducer and activator of transcription
TNF-α: Tumor necrosis factor alpha
TRPV4: Transient receptor potential vanilloid 4
VIM: Vimentin

---

**Supplementary Figure 1. Validation of Rat Immortalized Müller glia (rMC-1) (A)** Rat Immortalized Müller glia cultures (rMC-1) stained for canonical markers (vimentin [VIM], glutamine synthetase [GS], and SOX9) and negative markers (GFAP and IBA1). **(B)** LC_MS/MS analysis of polyunsaturated fatty acids (PUFAs) and lipoxygenase products released by immortalized Müller glia. Images acquired at 20x magnification; scale bar = 50 µm.

