## Supplementary figures and images for "Lipoxin B_4_ Mitigates TRPV4-Activated Müller Cell Gliosis During Ocular Hypertension"

### Supplemental Figure 1

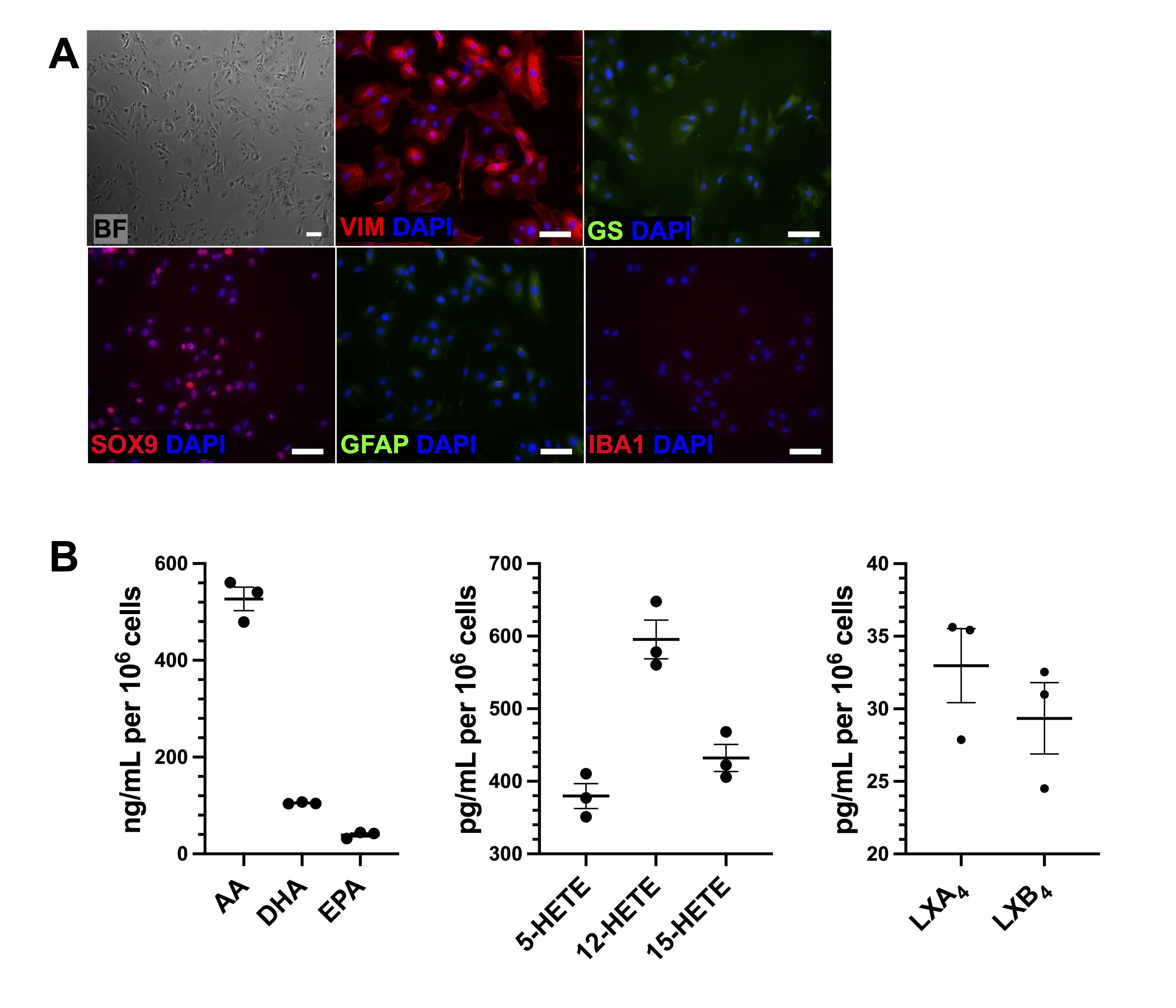
