## Supplementary material for "Lipoxin B_4_ Mitigates TRPV4-Activated Müller Cell Gliosis During Ocular Hypertension": Table 1

| **Primers Mouse** | **Sequence (5’ 🡪 3’)** |
| --- | --- |
| *Alox5* Forward | ACA GCT TAT CTG CGA GTA TGG |
| *Alox5* Reverse | GGG AAACAC AGG GAG GAA TAG |
| *Alox15* Forward | TGG GTT CTC TGC CTT AGT GG |
| *Alox15* Reverse | CAC TCA GGG TTG TCA CCT CA |
| *Il6* Forward | CCC CAA TTT CCA ATG CTC TCC A |
| *Il6* Reverse | CGC ACT AGG TTT GCC GAG TA |
| *TNF-α* Forward | TGA TCG GTC CCC AAA GGG AT |
| *TNF-α* Reverse | TGT CTT TGA GAT CCA TGC CGT |
| *Stat3* Forward | ACC AACGAC CTGCAG CAA TA |
| *Stat3* Reverse | TCC ATG TCA AAC GTG AGC GA |
