## Supplementary material for "Lipoxin B_4_ Mitigates TRPV4-Activated Müller Cell Gliosis During Ocular Hypertension": Table 2

| **Primers Rat** | **Sequence (5’ 🡪 3’)** |
| --- | --- |
| *Alox5* Forward | AGA GTC AAG AAT CTG GTG GGC |
| *Alox5* Reverse | GGT GAC AGT GTA GGA AGG CA |
| *Alox15* Forward | GGG ACT CGG AAG CAG AAT TCA A |
| *Alox15* Reverse | GCC CTG AAC CCA TCG GTA A |
| *Il6* Forward | TCC TAC CCC AAC TTC CAA TGC TC |
| *Il6* Reverse | TTG GAT GGT CTT GGT CCT TAG CC |
